# Neuronal hyperactivity in a LRRK2-G2019S cellular model of Parkinson’s Disease

**DOI:** 10.1101/2021.06.23.449591

**Authors:** Edinson Lucumi Moreno, Siham Hachi, Sarah L. Nickels, Khalid I.W. Kane, Masha Moein, Jens C. Schwamborn, Alexander Skupin, Pieter Vanden Berghe, Ronan M.T. Fleming

## Abstract

Monogenic Parkinson’s Disease can be caused by a mutation in the leucine-rich repeat kinase 2 (LRRK2) gene, causing a late-onset autosomal dominant inherited form of Parkinson’s Disease. The function of the LRRK2 gene is incompletely understood, but several *in vitro* studies have reported that LRRK2-G2019S mutations affect neurite branching, calcium homeostasis and mitochondrial function, but thus far, there have been no reports of effects on electrophysiological activity. We assessed the neuronal activity of induced pluripotent stem cell derived neurons from Parkinson’s Disease patients with LRRK2-G2019S mutations and isogenic controls. Neuronal activity of spontaneously firing neuronal populations was recorded with a fluorescent calcium-sensitive dye (Fluo-4) and analysed with a novel image analysis pipeline that combined semi-automated neuronal segmentation and quantification of calcium transient properties. Compared with controls, LRRK2-G2019S mutants have shortened inter-spike intervals and an increased rate of spontaneous calcium transient induction.

## Introduction

Parkinson’s disease (PD) is the second most common neurodegenerative disease affecting more than 10 million people worldwide [30, 1]. It is known that cell death in PD occurs at many levels of the nervous system [30], including the loss of cholinergic neurons in the pedunculopontine nucleus, noradrenergic neurons in the locus coeruleus and dopaminergic neurons from the substantia nigra pars compacta (SNc) [55, 3]. Dopaminergic neurons produce the neurotransmitter dopamine, which plays a central role in brain function [12, 52, 41]. Moreover, the loss of dopaminergic neurons is responsible for the primary motor symptoms in PD, including rigidity, resting tremors, bradykinesia and postural instability [35]. Further understanding of PD at the cellular level requires advanced patient-derived cellular models that recapitulate the main characteristics of the neurons that are selectively vulnerable to degeneration in PD.

Between 5-10% of PD cases have a genetic or familial origin [29, 49]. Certain mutations in the leucine rich repeat kinase 2 (LRRK2) gene are associated with familial and sporadic forms of PD [25]. LRRK2 is a multi-domain protein localised in the cytoplasm. It has kinase and GTPase activity in addition to domains for protein-protein interactions that make it a key regulator of cellular function [16, 39, 15, 50]. However, its exact function is still incompletely understood. Mutation G2019S is the most common LRRK2 mutation, it has been detected in 1% to 3% of sporadic PD cases and 3% to 6% in familial PD cases world-wide [18, 27]. In addition, clinical symptoms associated with patients with G2019S mutation include tremor, dystonia, cognitive impairment/dementia and anxiety [24, 27]. Mutant LRRK2 proteins impair normal protein phosphorylation in DA neurons and affect cell survival [18]. Over-expression of LRRK2 results in mitochondrial fragmentation [62] and an increase in LRRK2 activity is involved in neuronal apoptosis via mitochondria. Furthermore, LRRK2 deletion protects against mitochondrial dysfunction [16, 23] and a role for this protein in cytoskeletal dynamics has been demonstrated [18, 32, 6]. Several studies reported that LRRK2 mutations alters neuronal morphology, manifested by reduced neurite length as well as complexity [32, 10]. However, little is known about the effect of LRRK2 mutations on neuronal activity.

The study of PD at the cellular level has been facilitated by the use of induced pluripotent stem cell (iPSCs) technology. Several methods have been developed to reprogram human somatic cells into iPSCs. Reinhardt et al. [48] demonstrated the use of iPSC technology to obtain human neuroepithelial stem cells (hNESC) using small molecules. These hNESC can then be differentiated into specific neuronal cell types, including midbrain dopaminergic neurons (mDNs). Monogenic patient-derived stem cells can also be genetically modified to repair certain mutations, resulting in isogenic control lines, with the same genetic background as the patient-derived cells. Dopaminergic neurons derived from patient’s iPSC with LRRK2 mutation G2019S have shown increased expression and accumulation of alpha synuclein protein, shortening of neurite length together with diminished branching [22], increased susceptibility to oxidative stress [42], mitochondria damage [51] and signs of neurodegeneration in general [51, 22].

In experimental neuroscience there are several options to record electrophysiological activity of neurons *in vitro.* Patch clamp recording techniques provide very accurate measurements, however, they cannot be used to measure the electrophysiological activity of a large population of neurons. An alternative technique is to use various fluorescent indicators of electrophysiological activity, coupled with light microscopy and digital cameras [14]. Fluorescent calcium indicators respond to the binding of calcium ions by changing their fluorescence properties. *Calcium imaging* is the combination of fluorescent calcium indicators with imaging instrumentation, and is a versatile technique for studying neuronal activity and different aspects of cellular development [28, 36]. In particular, fluorescent calcium-sensitive dyes [56] permit simultaneous recording of electrophysiological activity in a large population of neurons. *Calcium imaging* has found widespread applications in neuroscience and its ability to record electrophysiological activity for a large population of neurons simultaneously makes it especially suited to analysing the phenotypic differences between normal and diseased neurons.

Neuronal activity is characterised by firing single or bursts of action potentials. During an action potential, intracellular calcium concentration increases transiently [54, 26]. A calcium transient is characterised by a fast rise in intracellular calcium concentration followed by a multi-exponential decay. Calcium imaging is an accessible option to characterise the neuronal activity of differentiated neurons derived from patient samples [31]. This approach is based on the use of fluorescent indicators that are sensitive to calcium concentration, using either calcium sensitive dyes (e.g. Fura-2 AM and Fluo-4 AM) [4] or genetically encoded calcium sensitive protein sensors [2]. With a suitable microscope and camera, one then obtains a temporal series of fluorescent images.

Calcium imaging can generate a large amount of data, necessitating the development of new computational techniques for processing and analysis to leverage the potential of this technique [47]. Given a temporal series of fluorescent images, individual neurons can be identified by manually selecting regions of interest. However, this method is subjective and laborious when large populations of neurons are being imaged. Therefore, robust and reliable computational approaches are required to automate and facilitate the analysis of calcium imaging data [44, 45]. Automated identification of individual cells is one of the main image analysis challenges. Different methods exist to identify cells, including machine learning [57], non-negative matrix factorisation [34], independent component analysis [40] and sparse dictionary learning [17]. Most of these approaches typically delineate regions of interest corresponding to cell bodies and then generate fluorescence traces for each cell. As these traces represent the dynamics of calcium indicators, which are much slower than the spiking activity of neurons, various algorithms have been developed to deconvolve a fluorescence signal into a spike train [65]. Such methods are based on Markov chain Monte Carlo [60], fast non-negative deconvolution [59], greedy algorithms [20] or finite rate of innovation[43].

Combining these methods together in one pipeline would be useful for the neuroscience community. Patel et al. [44] developed a pipeline to measure features of neuronal network activity. It uses a cell segmentation method based on Independent Component Analysis that provides accurate results. However, it is semi-automated and cannot identify cells that are not active or cells firing synchronously. Giovannucci et al. [19] developed automatic and scalable analysis methods to address problems common to preprocessing, including motion correction, neural activity identification, and registration across different sessions of data collection. Cantu et al [8] developed a graphical user interface-based and automated analysis software package for motion correction, segmentation, signal extraction, and deconvolution of calcium imaging data.

In this paper, we developed a fully automated pipeline for calcium image analysis and analysis of fluorescence traces from individual neurons. This pipeline was applied to quantitatively characterise the neuronal activity of human neuroepithelial stem cell-derived neurons *in vitro.* We used a sparse dictionary learning based-algorithm [17] to identify single neurons in a spontaneously firing population, enabling the extraction of fluorescence traces for individual neurons. Fluorescence traces were deconvolved into spike trains [59] and calcium transients were identified and extracted to quantify their properties and characterise the neuronal activity of differentiated cells. Applying this image analysis pipeline to calcium time-series of spontaneously firing iPSC-derived wild-type and LRRK2-G2019S mutant neurons, revealed neuronal hyperactivity in mutant neurons. Although the LRRK2-G2019S mutation is associated with neuronal hyperactivity in vitro, the causal mechanisms remain to be elucidated in future work.

## Results

### Automated neuronal segmentation of in vitro calcium time-series

A control human neuroepithelial stem cell-line (K7), and the same cell-line with an introduced LRRK2-G2019S mutation (K7M), in addition to a LRRK2-G2019S patient-derived cell-line (T4) and its isogenic control (T4GC), were maintained and differentiated into midbrain dopaminergic neurons (Figure 1). An automated analysis pipeline (Figure 5) was implemented for quantification of neuronal activity in calcium imaging data and applied it to spontaneously firing hNESC-derived neurons. An automated sparse dictionary learning algorithm was adapted and successfully applied to segment calcium imaging data (Figure 7a).

**Figure 1:**
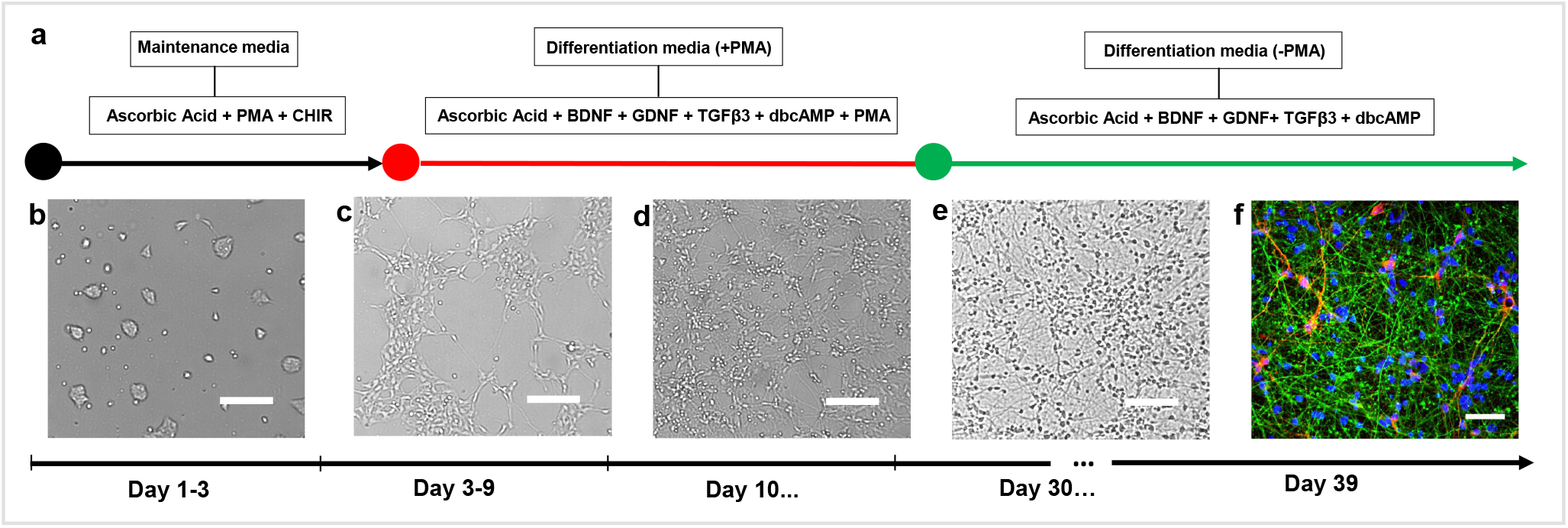
Differentiation of hNESC into dopaminergic neurons. (a) Protocol using small molecules. Bright field images of (b) hNESCs 1 day after seeding, cultured in maintenance media containing N2B27 medium, ascorbic acid, PMA and CHIR. (c) hNESCs 3 days after seeding, cultured in differentiation media containing N2B27 medium ascorbic acid, BDNF, GDNF, TGF 3, dbcAMP and PMA. (d) hNESCs 10 days after seeding, cultured in differentiation media as in (c) but without PMA. (e) hNESCs differentiated into neurons in a well of a 96 well plate after 1 month differentiation. (f) Immunostaining of selected region in the well, showing neurons positive for nuclei with Hoechst (blue), TUB III (green) and TH (red); scale bar 50 m.

An optimal wavelet scale parameter was identified (scale 4) as smaller wavelet scales yielded in-homogeneous fluorescence intensity within a single region of interest representing a neuron, leading to over-segmentation (Figure 2b scale 3). Larger wavelet scales exceeded the limit of the dimensions of our cells including other cells or parts of their neighbourhood resulting in erroneous segmentation (Figure 2c scale 5). The choice of the optimal scale parameter strongly depends on size of the cells being analysed (Figure 2b-c). The optimal pixel intensity threshold was determined to be 0.06, which represents the percentage of the maximum intensity of the smoothed image. After optimising the different parameters, individual neurons were accurately and automatically segmented and their corresponding fluorescence traces were analysed (Figure 7c). These traces revealed calcium transients with different waveforms and different frequencies showing the diversity in firing frequencies and firing patterns in this neuronal culture.

**Figure 2:**
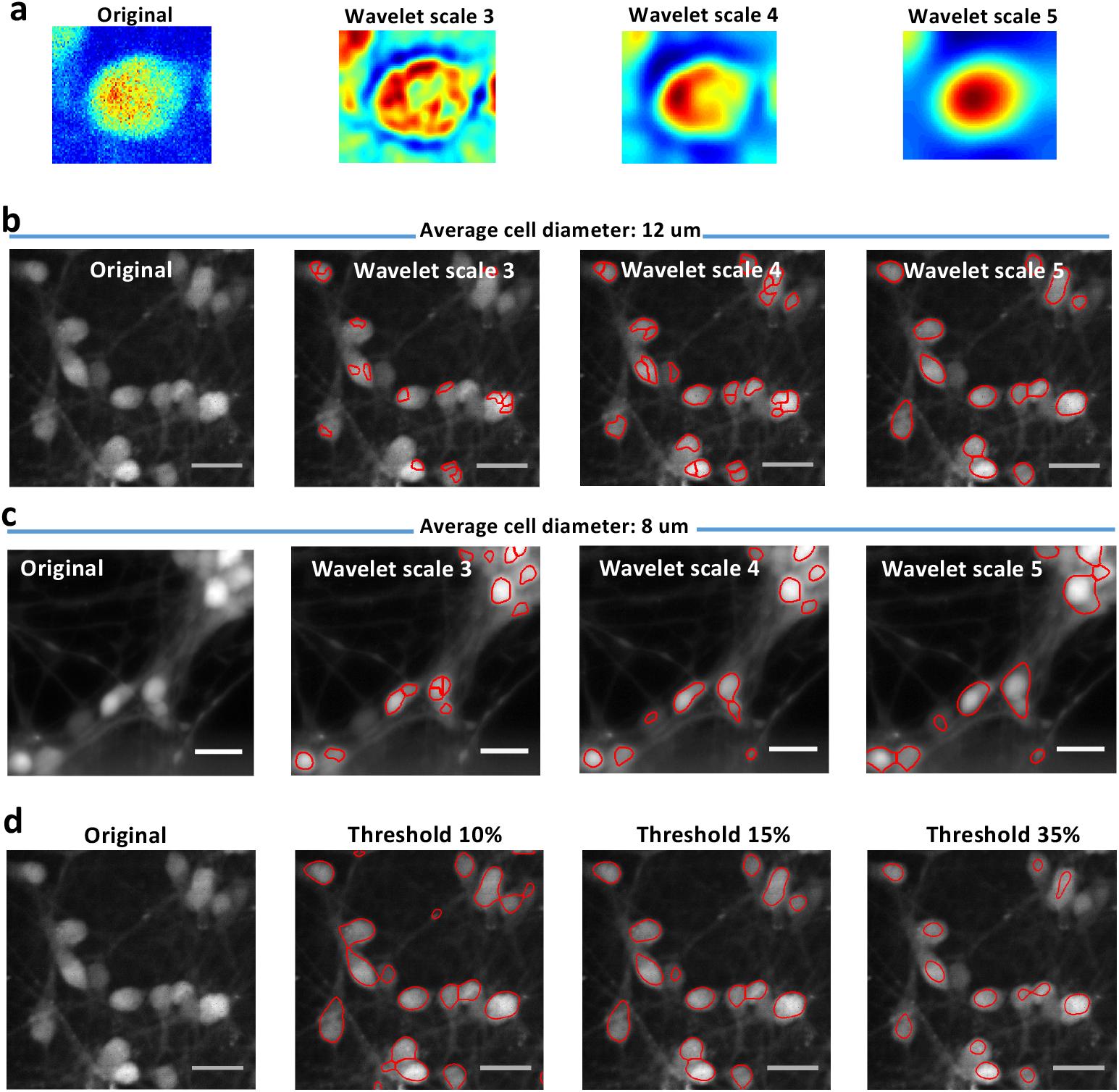
Effect of wavelet scale size on segmentation. **(a)** Different wavelet scale sizes have different effects on the image in the preprocessing step. The purpose of the wavelet function is to smooth the image and give more intensity to round shapes such as cell bodies, (b) The effect of different wavelet scales on segmentation of somas of an average diameter of 12 m. Clearly, wavelet scale 5, compared with 3 and 4, induces a more accurate segmentation as the somata are well delineated, **(c)** The effect of different wavelet scales on segmentation of somas of an average diameter of 8 m. Clearly, wavelet scale 4 performs better than 3, where some somata are subdivided into several parts and 5, where some somata are merged in one segment, **(d)** Effect of intensity threshold on segmentation.

### Hyperactivity of LRRK2-G2019S mutant neurons

After neuronal segmentation and fluorescence traces, with more than 5 calcium transients, were obtained from a large number of neurons for each cell-line (1258 ±336 traces for K7, f022±421 for K7M, 480±f95 for T4GC and 369 ± 267 for T4, *n =* 3 plates). For each biological replicate, inter-spike interval values of a pair of cell-lines (e.g. K7 vs. K7M) were merged together to form one set (Figure 3a). Each set was then classified according to short, medium and long classes of inter spike intervals (Figure 3b). Each class contained average inter spike interval from both control and mutant neurons (Figure 3c). Within the short inter-spike interval class (0 - 10 sec), the proportion of mutant neurons is significantly higher than for controls. The proportion of mutant neurons decreases as the inter-spike intervals increases. Approximately 55% of neurons with a LRRK2-G2019S mutation exhibited inter-spike intervals between 0-10 sec against 32%) in control (Figure 3d). Figure 3 shows data of one biological replicate for K7 and K7M cell-lines. The same analysis was performed on the other pair of cell-lines (T4 vs. T4GC) and for each biological replicate. No significant differences were observed in other calcium transient features (see Supplementary Material Figure 8 and 9). In summary, compared with isogenic controls, fluorescence traces from LRRK2-G2019S mutant neurons demonstrated reduced inter-spike intervals.

**Figure 3:**
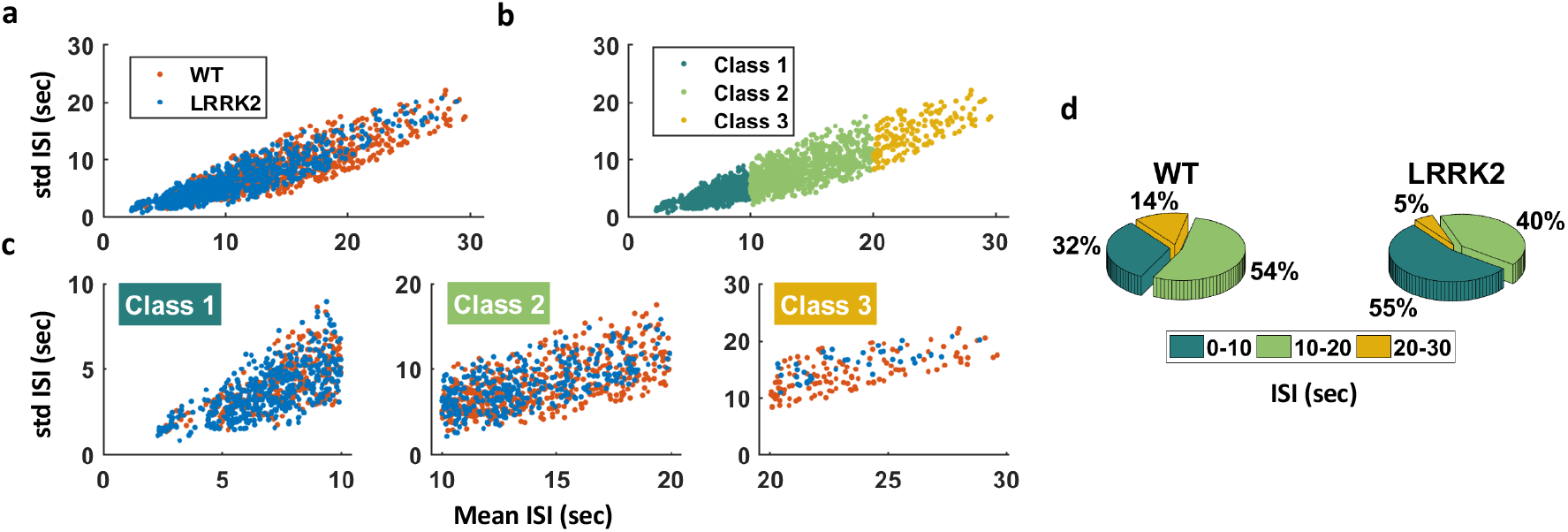
Classification of mean interspike interval (ISI) values (K7 and K7M cell-lines). **(a)** Scatter plot of merged inter-spike intervals (ISI, seconds) from control and LRRK2-G2019S mutants, **(b)** Classification of inter-spike intervals into 10-sec interval groups, colours correspond to individual classes (1-3). **(c)** Scatter plots of the merged inter-spike intervals from control and LRRK2-G2019S neurons for each class, (d) Pie charts showing relative proportions of neurons in each inter-spike interval class colour-coded same as in **(b).**

## Discussion

Calcium imaging provides a powerful means to monitor the activity of neuronal populations *in vivo* 64] and *in vitro* [21], In contrast to electrophysiological approaches, such as patch clamp techniques, it allows simultaneous recording of neuronal activity and spatial location of individual neurons for a large population [21], Calcium imaging generates a large amount of data, which becomes challenging to analyse manually as neuronal population size increases. It is therefore necessary to apply computational methods to partially automate the analysis of calcium imaging data.

Computational image analysis is required to efficiently, reliably and quantitatively analyse calcium time-series and activity-dependent fluorescence traces. With the recent developments of several algorithms to segment calcium imaging data, infer spike trains from fluorescence traces and analyse calcium transients [46], it is necessary to establish pipelines of methods as one package. In this study, we established a pipeline that brings together a set of complementary algorithms to analyse calcium imaging data and quantify calcium transient properties to characterise neuronal activity.

In this pipeline, we integrated a sparse dictionary learning-based segmentation algorithm [17] that was shown to be more robust (94.3% sensitivity) than the CellSort tool [40] (83.1% sensitivity). The sparse dictionary learning-based algorithm provides an accurate segmentation and can discriminate two overlapping cells exhibiting similar firing patterns. In the preprocessing step, this method uses a wavelet transform to intensify the fluorescence traces belonging mainly to cell bodies and not processes. This limits the amount of segmented regions such as neuronal projections that might not be of interest to the user. These wavelet functions are described by a convolution mask with the coefficients [1 4 6 4 1]/16. This function can be extended to different scales by adding zeros between the coefficients [47]. Choosing the right wavelet scale is crucial to obtain well segmented cells and depends on the size of the imaged cells (Figure 2).

In addition to the wavelet scale, other parameters need to be tuned for an optimal segmentation, such as the intensity threshold. If the threshold is too low, then regions containing pixels with a low intensity that do not belong to a cell body would be considered. On the other hand, if the threshold is too high then cell bodies with lower intensity pixels would be excluded. The requirement for tuning these parameters makes the pipeline semi-automated, when considering all cell types, and fully automated when one cell type is being considered. Tuning these parameters strongly depends on the experimental setup used for calcium imaging recordings, such as type of microscope, camera, fluorescence light source and calcium indicators. However, once these parameters are optimised for a particular calcium imaging acquisition protocol, automated processing of similar datasets is possible. Although there is a significant parameter tuning in this approach, it provides robust segmentation of individual neurons. This pipeline was also effectively applied to different cell culture systems, such as organoids [38] and microfluidic cell culture [31].

Patient-derived neurons are becoming extensively used in neuroscience for modelling and investigating different neurodegenerative diseases, revealing various relevant disease mechanisms [63]. We applied our image analysis pipeline to human neuroepithelial stem cells differentiated into dopaminergic neurons to measure the effect of a LRRK2-G2019S mutation on neuronal activity. Mutant LRRK2 has been found to be implicated in several cellular dysfunctions including mitochondrial degradation [11], calcium homeostasis impairment [11], synaptic transmission [13] as well as neurite length shortening [32].

We found that LRRK2-G2019S mutated neurons displayed a form of neuronal hyperactivity compared with control neurons. In particular, we demonstrated that the inter-spike interval between action potential-evoked calcium transients was significantly shorter in LRRK2-G2019S mutated neurons. As yet, we do not know the mechanism for this hyperactivity. However, this phenotype has been previously observed in several studies related to neurodegeneration in Alzheimer’s disease [7]. In this respect, an *in vivo* whole-cell patch clamp technique was utilised to assess the electrical activity of neurons vulnerable in Alzheimer’s disease [53]. This study aimed at linking the dendritic morphological abnormalities, in particular shortening of dendritic length, to neuronal excitability. A full morphological computational model, simulating *in vivo* conditions, predicted high firing frequency subsequent to reduction of dendritic length, making the neuron electrically more compact, thus leading to shorter inter-spike intervals, which was confirmed by patch-clamp recordings [53]. Siskova et al. reported that in a neuron with shortened morphology, synaptic currents would translated faster into postsynaptic currents resulting in hyperexcitability. This structure-dependant cellular mechanism could be a possible explanation to the hyperexcitability of LRRK2-G2019S mutants that we observed, considering that this mutation induces dendritic shortening [11]. In fact, Cherra et al. reported that the LRRK2-G2019S mutation decreases the dendritic mitochondrial density thus leading to dendritic area reduction [11]. Moreover we previously showed significantly shortened morphology of hNESC-derived neurons carrying a LRRK2-G2019S mutation [5].

Hyperactivity was previously observed in mouse and patient models of Alzheimer’s disease [7, 58]. Moreover, patient-derived motor neurons were shown to exhibit a form of neuronal hyperexcitability in amyotrophic lateral sclerosis disease [61] as well as cortical neurons in Huntington’s disease [9]. It is an interesting question as to why hyperactivity of vulnerable neurons seems to be a common pathological phenotype in these neurodegenerative diseases.

We combined different computational data analysis approaches into one pipeline to analyse calcium imaging data and calcium transients with the aim of quantifying neuronal activity of large neuronal ensembles. The pipeline includes accurate cell segmentation, fluorescence trace measurement, spike train inference and calcium transient analysis. This pipeline will facilitate investigations of neuronal activity with large numbers of neurons. We assert that new imaging techniques, computational approaches and induced pluripotent stem cell technology taken together have great potential to facilitate identification of disease phenotypes, discovery of potential biomarkers and novel therapeutic strategies.

In conclusion, we successfully applied a calcium imaging pipeline to human neuroepithelial stem cells with a LRRK2-G2019S mutation that were differentiated into dopaminergic neurons *in vitro* and compared them to calcium imaging of isogenic controls. We observed evidence for a disease-specific phenotype characterised by neuronal hyperactivity, similar to what has been reported in the most common neurodegenerative diseases such as Alzheimer’s disease and amyotrophic lateral sclerosis. Further studies on neuronal hyperexcitability in LRRK2 mutated neurons can be expanded to investigate whether this phenotype is manifested in other LRRK2 mutations (e.g. R1441C mutation) and whether it persists as neurons mature.

## Experimental procedures

### hNESC differentiation into dopaminergic neurons

A control human neuroepithelial stem cell-line (K7), and the same cell-line with an introduced LRRK2-G2019S mutation (K7M), in addition to a LRRK2-G2019S patient-derived cell-line (T4) and its isogenic control (T4GC), were maintained and differentiated into midbrain dopaminergic neurons using small molecules and growth factors, in 96 well plates, by using an existing protocol described by Reinhardt et al. [48]. Here we used 3 plates representing 3 biological replicates. In brief, CellCarrier optically clear bottom, black, tissue treated 96 well plates (Perkin Elmer) were coated with Matrigel (catalogue number 354277, lot number 3318549, Discovery Labware, Inc., Two Oak Park, Bedford, MA, USA). At the time of cell seeding, the coating medium was removed from each well of the 96 well plate and the different cell-lines were seeded in selected wells, at a density of 0.3 million cells/ml in 80 L of medium, and incubated at *37°C* with 5% CO_2_. The medium was changed every other day and this process was repeated throughout the differentiation process for 5 weeks.

The culture medium preparation “*N2B27 medium*” consisted of mixing equal amounts of Neurobasal medium (Invitrogen/life technologies) and DMEM/F12 medium (Invitrogen/life technologies) supplemented with 1% penicillin/streptomycin (life technologies), 2 mM L-glutamine (life technologies), 0.5 X B27 supplement without Vitamin A (life technologies) and 0.5X N2 supplement (life technologies).

The medium to maintain the hNESC in culture *“maintenance medium”* consisted of N2B27 medium with 0.5 M Purmophamine (PMA) (Enzo life sciences), 3 M CHIR (Axon Medchem) and 150 M Ascorbic Acid (Sigma Aldrich). The differentiation medium formulation to induce the differentiation of hNESC towards midbrain dopaminergic neurons *“differentiation medium with PMA”* consisted in N2B27 medium with 200 M ascorbic acid, 0.01 ng/ L BDNF (Peprotech), 0.01 ng/ L GDNF (Peprotech), 0.001 ng/ L TGF 3 (Peprotech), 2.5 M dbcAMP (Sigma Aldrich) and 1 M PMA. 80 L of differentiation medium with PMA was changed every 2 days during the first 6 days of culture in the differentiation process. For the maturation of differentiated neurons PMA was no longer added to the differentiation medium *“differentiation medium without PMA”* from day 7 onwards, this differentiation medium without PMA was changed every 2 days during 3 weeks.

### Calcium imaging

A calcium imaging assay was done on representative wells of a black CellCarrier optically clear bottom, tissue treated 96 well-plate (Perkin Elmer). Differentiation medium was removed from the wells with differentiated cells. Those wells were washed twice with 100 L of neurobasal medium. Afterwards, the washing medium was removed and 80 L of 5 M cell permeant Fluo-4 AM (Life technologies) in neurobasal medium was added to wells at room temperature and incubated at 37 °C for 30 minutes. The medium with Fluo-4 was removed and replaced by the old differentiation medium and left incubating for 10 more minutes. Images of spontaneously firing hNESC-derived neurons were acquired, using an epifluorescence microscope (Leica DMI6000 B) equipped with a confocal device (Revolution DSD2, Andor) and an sCMOS camera (Neo 5.5, Andor). Images of size 1200 x 1200 pixels were sampled at a rate of 2 Hz for 4 min, stored as separate image files and analysed using custom Matlab (version 2017b; MathWorks) scripts. Four to six calcium time-series of randomly selected fields of view were acquired in three representative wells for each cell-line.

### Automated image analysis

Calcium image analysis was achieved by combining published open source algorithms developed in MATLAB into a novel automated pipeline. The pipeline is illustrated in Figure 5 and consists of the following steps: 1) neuronal segmentation, 2) fluorescence trace extraction from the segmented neurons, 3) spike train inference and 4) calcium spike analysis. We integrated all of these steps in a pipeline to quantify neuronal activity. Calcium imaging datasets for each cell-line were stored in one folder to perform batch analysis, where all datasets were automatically analysed with one script.

**Figure 4:**
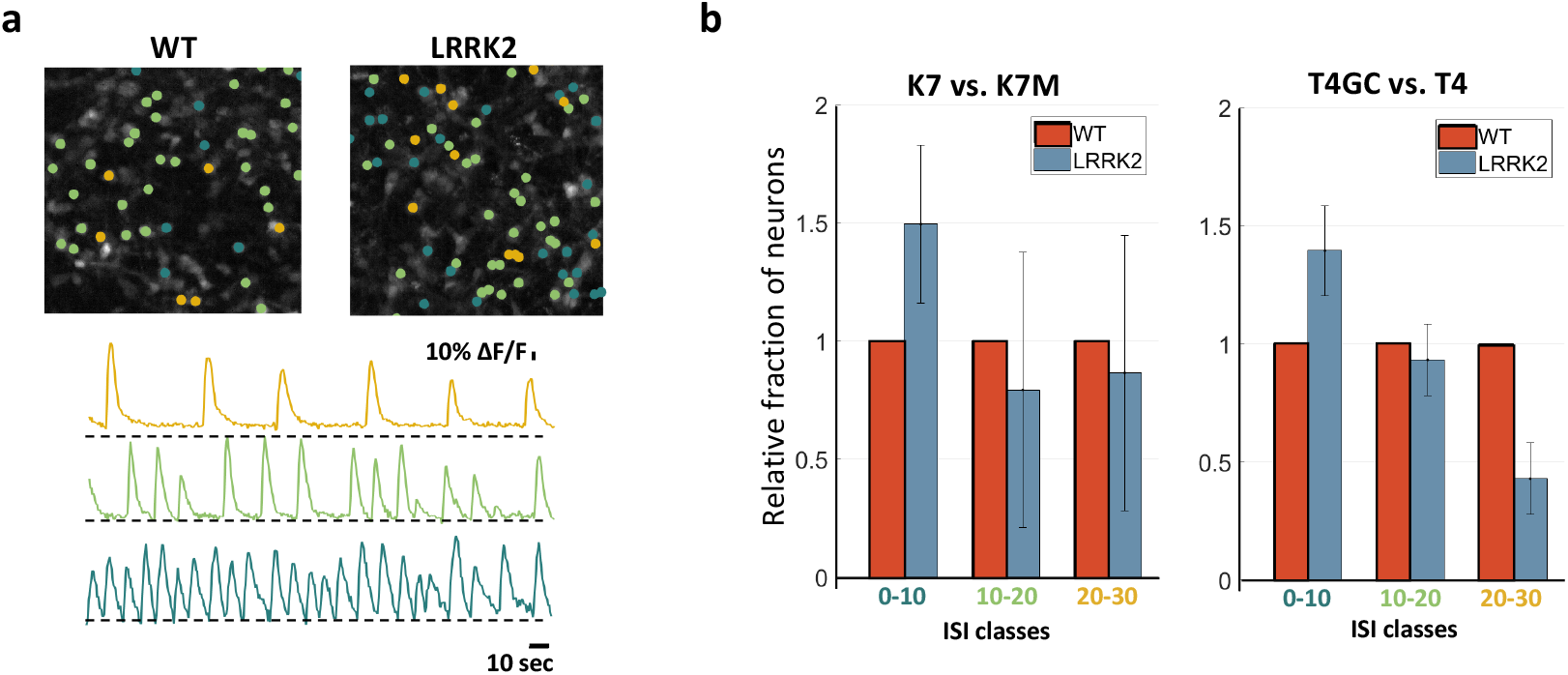
Hyperactivity of LRRK2-G2019S mutant neurons. **(a)** Mean fluorescence frames of K7 (control) and K7M (LRRK2-G2019S) cell-lines overlaid with activity maps (top) and examples of traces from each class (bottom), **(b)** Comparison of the fractions of LRRK2-G2019S neurons in each class relative to wild-type (for both cell-lines).

**Figure 5:**
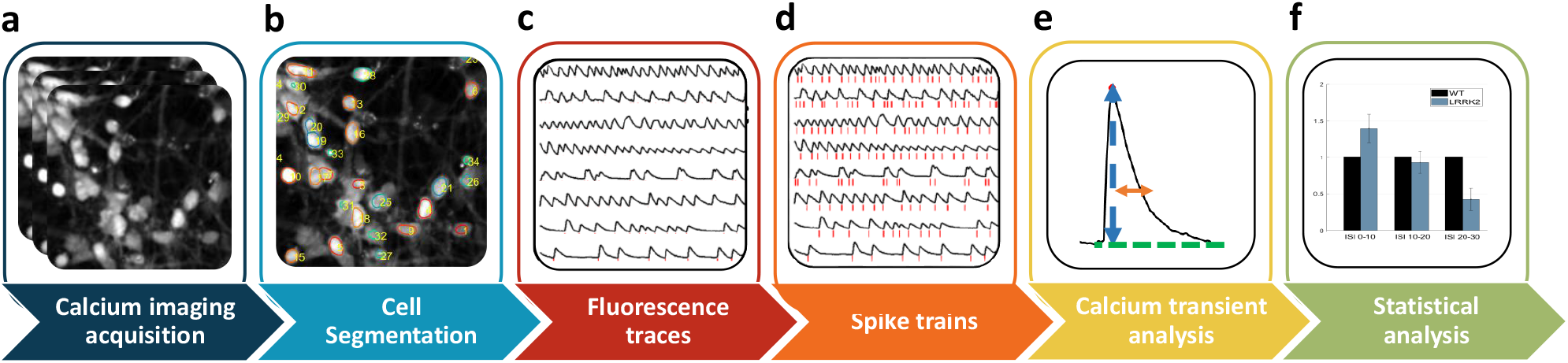
Automated calcium image analysis. Automated image analysis pipeline for calcium imaging data to quantify neuronal activity. Cells are cultured in an automated cell culture observatory **(a)**. Calcium imaging data of a neuronal population are acquired **(b)**. Regions of interest (ROIs) corresponding to individual neurons are identified **(c)**. Fluorescence traces (black traces) and their underlying spike trains (red lines) are measured for each ROI **(d)** Spike trains (red lines) are identified. **(e)**. Calcium transients are detected and their properties are estimated, as shown below in Figure 6 **(f)**. Statistical analysis is performed to estimate differences in calcium transient parameters between two different neuronal populations.

#### IMAGE SEGMENTATION

To detect individual neurons in calcium time-series, we employed an automated image segmentation technique, based on sparse dictionary learning [17]. This method was proven to outperform state-of-the-art cell segmentation algorithms in calcium imaging data, particularly the most commonly used method based on independent component analysis and principal component analysis (the CellSort tool) [40].

Briefly, the algorithm reshapes the (2+1)D calcium imaging data into a 2D matrix, then applies a matrix factorisation based on sparse dictionary learning [33] to decompose the data into a spatial and temporal components. This is followed by an image segmentation step where a wavelet transform and watershed algorithm are used to detect single cells. The wavelet functions are based on a cubic B-spline, to remove background noise and enhance fluorescence signals coming from the pixels that belong to neuronal cell bodies over other shapes [47]. The wavelet functions use a convolution mask with the coefficients [1 6 4 1]/16. This function can be extended to different scales and can therefore be applicable to different cell sizes. For a more detailed description of the wavelet functions refer to Reichinnek et al. [47]. Here, we used a wavelet scale of 4 (see Discussion).

#### Fluorescence trace generation and spike train inference

Fluorescence traces were extracted from automatically segmented regions of interest, corresponding to neuronal somata, by averaging the intensities of all pixels within each region of interest at each frame. These traces are presented as relative changes in fluorescence intensity (Δ*F/F = (f - f*_0_)/*f*_0_) where *f* is the fluorescence intensity and *f*_0_ the fluorescence baseline (Figure 7) defined as the 20^th^ percentile of the fluorescence signal within a sliding window of 10 frames. Thereafter, the user can choose between spike train inference or peak detection corresponding to calcium transient peaks. In the case of spike train inference, fluorescence traces are deconvolved into spike trains using a fast non-negative deconvolution (OOPSI) algorithm described in Vogelstein et al. [59] or the constrained-OOPSI [46]. The output of the OOPSI algorithm is the probability that a spike occurred at a given time frame. We set a probability threshold of 0.85 to exclude improbable spikes.

#### Calcium transient analysis

After measuring fluorescence traces, we quantified the features of calcium transients using the CaSiAn tool; a JAVA-based application for calcium signal analysis [37], which was integrated into our pipeline. First, transient peaks were detected by finding the local maxima of each fluorescence trace. We used a peak threshold of 20% of the maximum transient amplitude. Subsequently, for each identified calcium transient, a set of features were quantified. The features quantified were the inter-spike interval (ISI), that is the time between two transient peaks, spike amplitude, spike width, area under the spike, calcium increasing rate and decreasing rate related to calcium fluxes (cf Figure 6). Fluorescence traces with less than 5 spikes were excluded.

**Figure 6:**
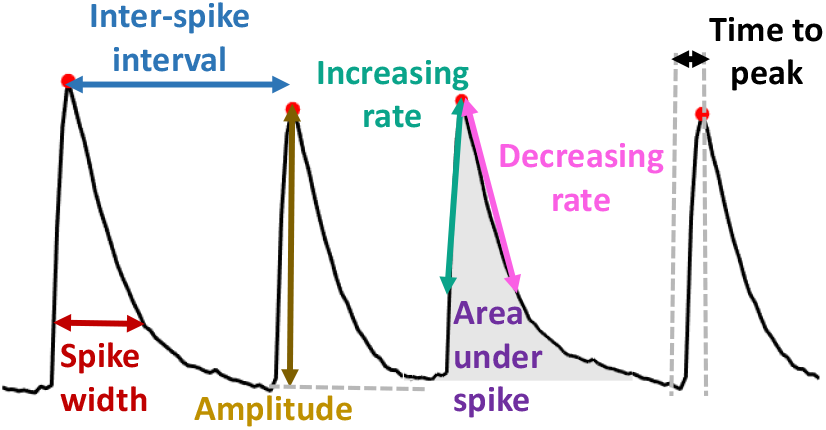
Description of the measured calcium transient features.

**Figure 7:**
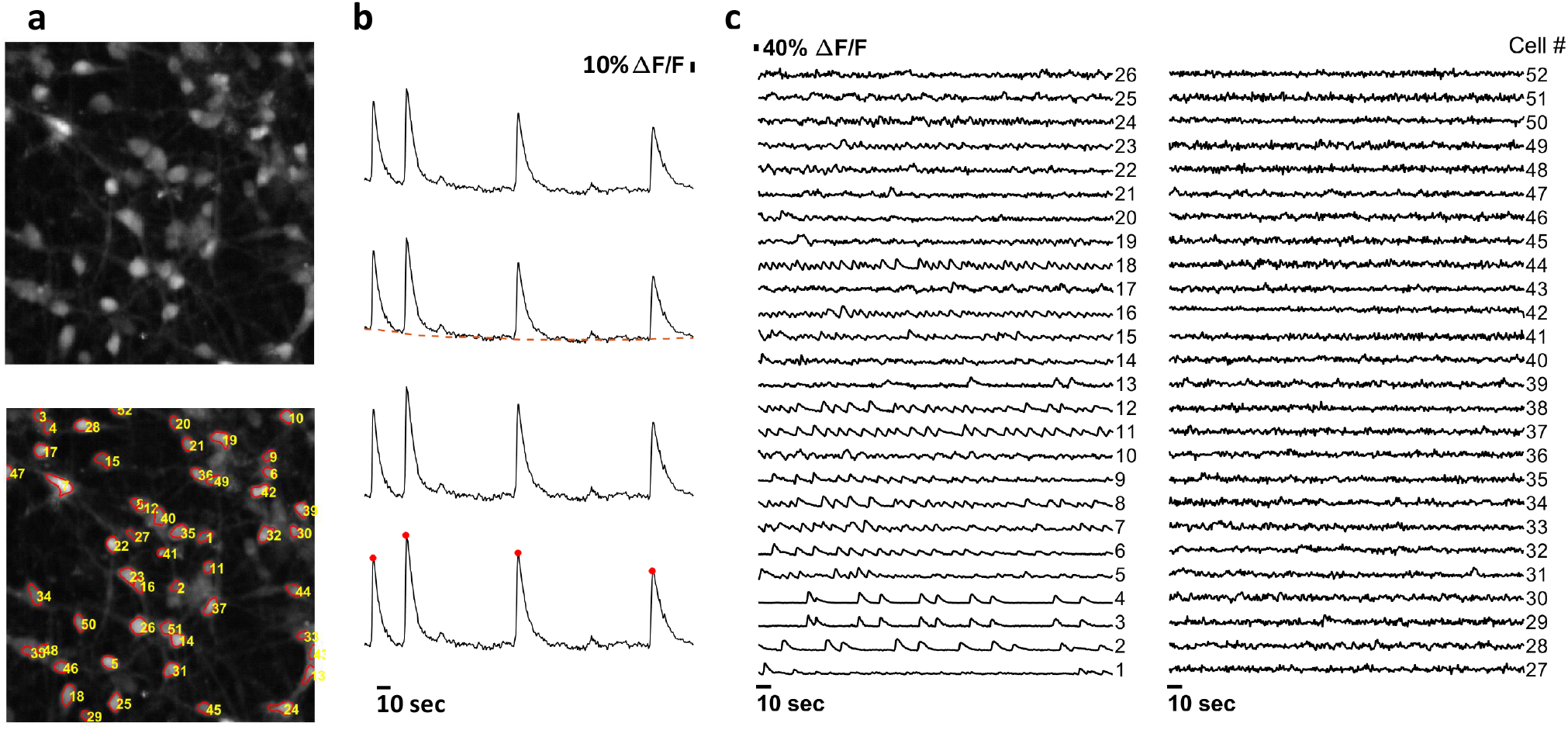
Automated segmentation, fluorescence trace estimation and spike train inference. Automated segmentation, fluorescence trace estimation and spike train inference of calcium time-series of a neuronal population. **(a)** Mean fluorescence frame of a calcium time-series (top) and numbered segmentation of individual neuronal somata (bottom). **(b)** Fluorescence baseline correction. **(c)** Fluorescence traces corresponding to the segmented neurons in **(a)**.

#### Comparison between LRRK2 G2019S mutants and isogenic control cell-lines

For each pair of mutant and isogenic control cell lines, the quantified features of calcium transients from sets of fluorescence traces from individual neurons were compared. The fluorescence traces from neurons of the same cell-line in different wells but the same biological replicate, were considered as a single set and subsequently the mean and standard deviation of each quantified feature from that set was estimated. Due to high variability in firing patterns, each fluorescence trace was classified according to its mean inter-spike interval (ISI), using intervals of 10 seconds, resulting in three classes (0 - 10, 10 - 20, 20 - 30 seconds). This classification was performed on each biological replicate of each cell-line. Statistical analysis of calcium transient features to calculate differences between wild-type and LRRK2-G2019S-mutated cell-lines. Specifically, a student’s t-test was used and P < 0.05 was considered statistically significant.

## Data and code availability

Control and LRRK2 patient derived raw calcium imaging data is accessible via the University of Luxembourg and subject to the EU General Data Protection Regulation 2016/679. Code may be accessed via the following version controlled repository https://github.com/rmtfleming/calciumPipe.

## Acknowledgements

This project has received funding from the European Union’s Horizon 2020 research and innovation programme under grant agreement No 668738. SH and ELM were supported by an Aides a la FormationRecherche training allowance from Fonds National de la Recherche Luxembourg (ref. 10099424). JCS lab is supported by a CORE grant from the Fonds National de la Recherche (ref. C13/BM/5791363), AS was supported by the Fonds National de la Recherche Luxembourg (ref. INTER/DFG/17/11583046).

## Author contributions section

Edinson Lucumi Moreno: Investigation, Methodology, Writing original draft, Writing review & editing, Project administration. Siham Hachi: Formal Analysis, Visualization, Writing original draft, Writing review & editing. Sarah L. Nickels: Investigation, Writing review & editing. Khalid I.W. Kane: Investigation, Writing review & editing. Masha Moein: Methodology, Writing review & editing. Jens C. Schwamborn: Funding acquisition, Supervision, Methodology, Resources, Writing review & editing. Alex Skupin: Funding acquisition, Supervision, Resources, Writing review & editing. Pieter Vanden Berghe: Conceptualization, Funding acquisition, Methodology, Resources, Supervision. Ronan M.T. Fleming: Conceptualization, Funding acquisition, Methodology, Resources, Writing original draft, Writing review & editing.

## Declaration of interests

The authors declare that the research was conducted in the absence of any commercial or financial relationships that could be construed as a potential conflict of interest.

## Supplementary material

Additional calcium transient features of LRRK2-G2019S mutants and controls for two cell lines, Figures 8 and 9.

**Figure 8:**
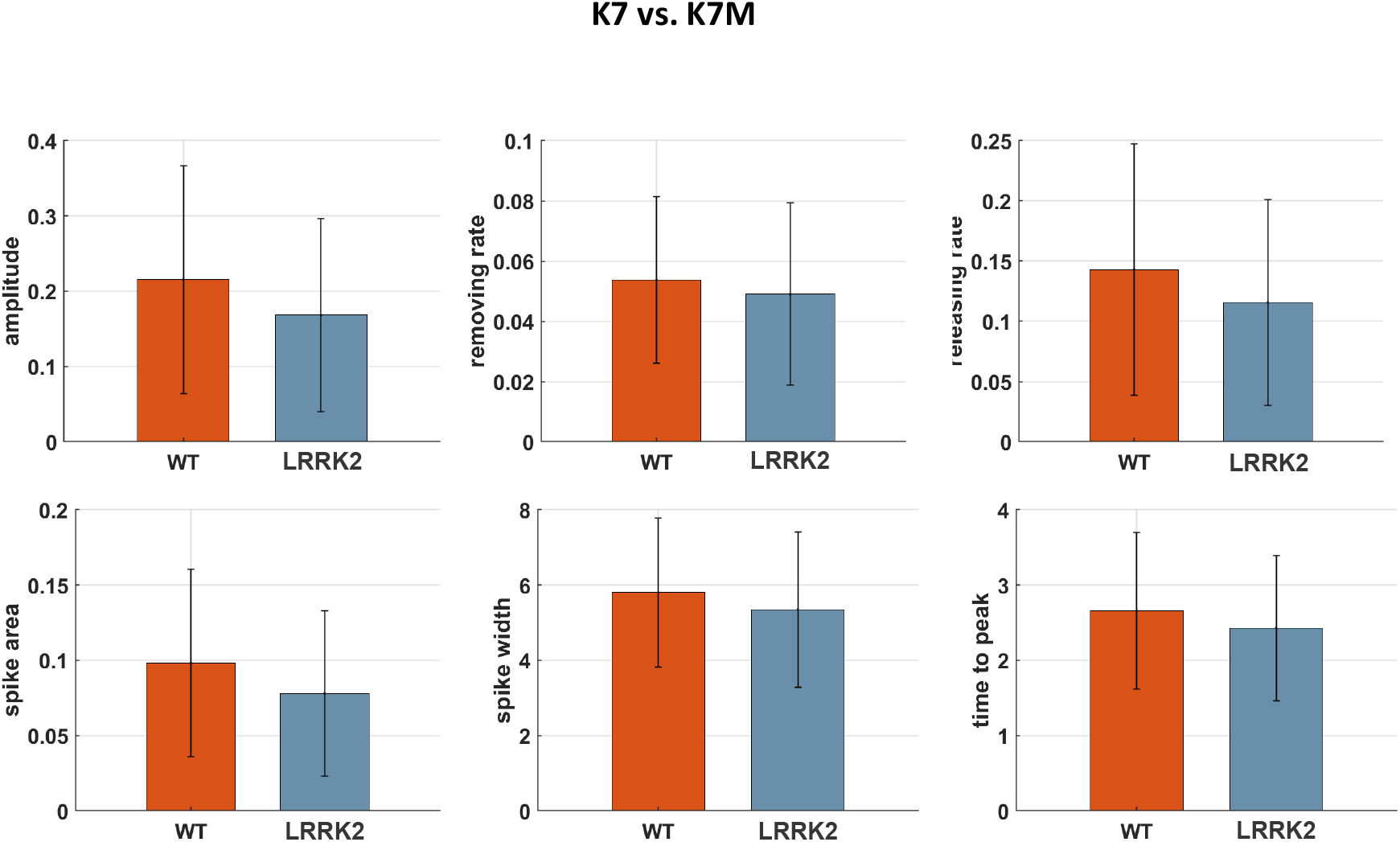
Additional calcium transient features of LRRK2-G2019S mutants and controls (K7 and K7M cell lines). No significant differences were observed.

**Figure 9:**
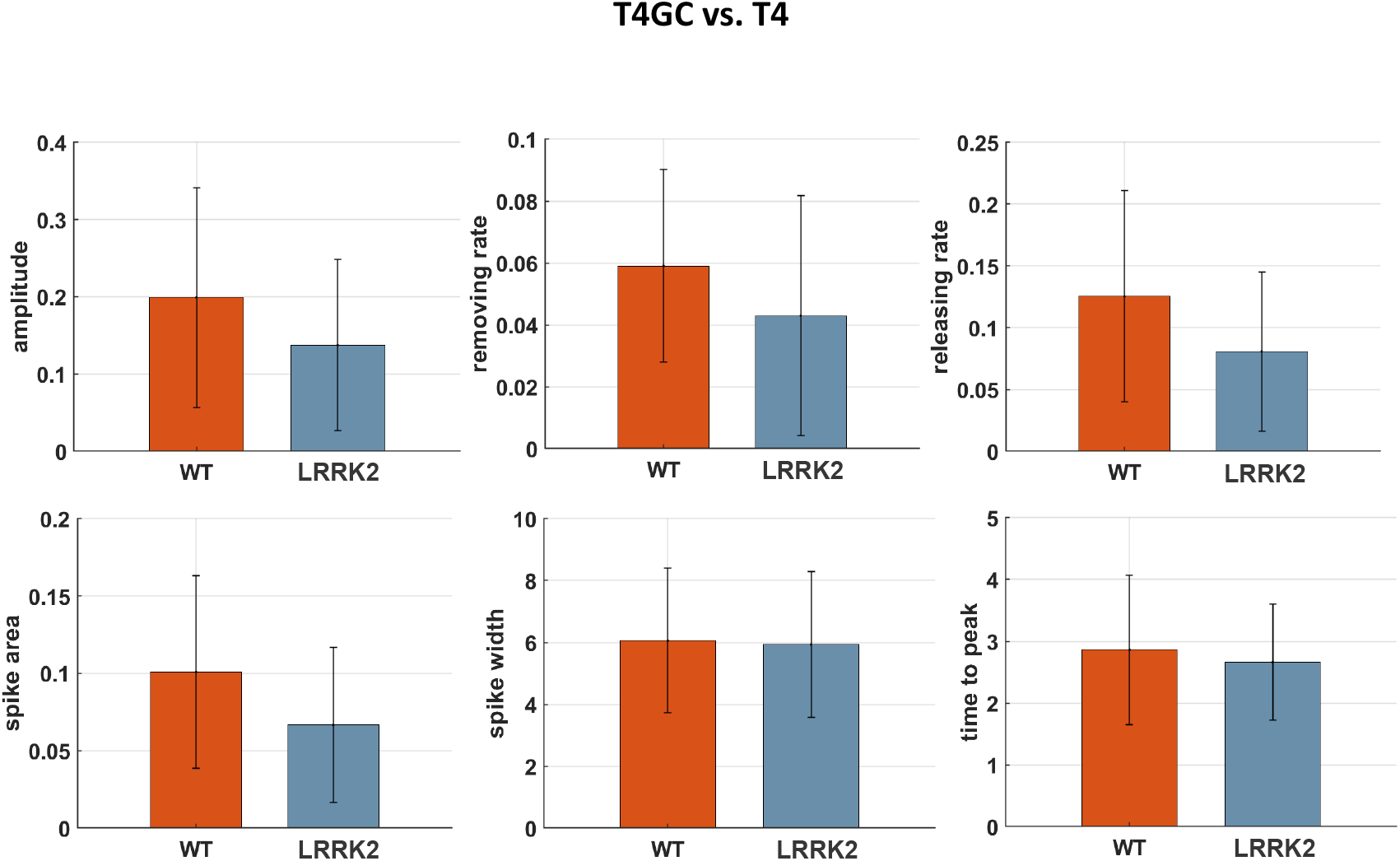
Additional calcium transient features of LRRK2-G2019S mutants and controls, T4 and T4GC cell lines respectively. No significant differences were observed.

## Notes

### Competing Interest Statement

The authors have declared no competing interest.

